# Tactile stimulation restores inhibited stretch reflex attributable to attenuation of la afferents during surprise landing

**DOI:** 10.1101/2022.09.14.507870

**Authors:** Yu Konishi, Ryo Yoshii, Daisuke Takeshita

## Abstract

**Purpose:** Arthrogenic muscle inhibition(AMI) is induced by pathological knee conditions. The present study aimed to investigate the effect of tactile stimulation on reflex changes induced by simulated AMI during unpredictable landing performances.

**Methods:** Twenty participants performed six unilateral landing tasks: 15cm normal landing (15NL), 30cm normal landing(30NL), surprise landing(SL), 30cm normal landing following vibration (30NLV), surprise landing following vibration(SLV), and surprise landing following vibration with Kinesiology-tape(SLK). For the surprise landing tasks, the solid landing platform(15cm) was removed and replaced by a false plate platform. Since the false plate dislodged easily under load, participants unpredictably fell through the plate to the actual landing surface 15cm below. After completing 15NL, 30NL, SL, vibration stimulation was applied to participants’ knees to induce neurological changes similar to AMI. After vibration stimulation, participants performed 30NLV, SLV and SLK in a random order. EMG in the post-landing short latency(31–60 ms) and medium latency(61–90 ms) periods were examined. EMG signals from the vastus lateralis(VL), vastus medialis(VM), and lateral hamstrings(LH) were recorded and compared between tasks.

**Results:** EMG of all muscles were significantly enhanced in the medium latency period. Enhanced EMG were suppressed by vibration stimulation in the VL, but the suppressed EMG were restored after cutaneous stimulation(p<0.01).

**Conclusion:** Our findings suggest that AMI could alter motor control patterns during unpredictable landing, and that tactile stimulation could restore the altered motor control to a normal state. The use of tactile stimulation may help maintain normal motor control patterns in injured individuals.

## Introduction

Quadriceps activation failure resulting from neurophysiological abnormalities is typically observed following knee injury or associated pathology and remains long after other symptoms abate(1–3). Previous studies reported that quadriceps activation failure was observed in patients with knee ligament injury(2) or osteoarthritis(4), elderly patients hospitalized after falls, and even healthy elderly people(5, 3). Quadriceps activation failure prevents patients from attaining maximal strength, even though the quadriceps itself is not damaged at all. Recently, the term arthrogenic muscle inhibition (AMI) has been used to describe the impaired ability to voluntarily contract the quadriceps(6, 7). The mechanism of AMI can be partially explained by the hypothesis set forth by Konishi et al.(8–10): impaired afferent feedback that travels from the structures around the knee joint, such as ligaments, capsules, and skin, to gamma motor neurons attenuates Ia afferents(11), and because adequate Ia afferent activity is necessary to recruit high-threshold motor units, maximal voluntary activation of the quadriceps cannot be achieved(12–14). In other words, hindered recruitment of high-threshold motor units is considered a possible mechanism underlying AMI.

This hindrance to motor unit recruitment is also observed in healthy individuals with muscle weakness after application of prolonged vibration(14). Indeed, several studies have reported a decline in the activity of alpha motor neurons in healthy individuals upon prolonged vibration stimulation(2, 14, 4). Thus, previous studies used neurological changes in the quadriceps induced by prolonged vibration stimulation as a simulated model of AMI(15, 16). Those studies found that cutaneous stimulation such as tactile and electrical stimulation to the skin could compensate for deficits in both strength and EMG activity following the application of prolonged vibration(15, 16). Moreover, according to a previous animal study that demonstrated the mechanism of how cutaneous stimulation affected motor function, sensory input from the skin around the joint could activate gamma motor neurons to modulate Ia afferent activity(11). Since Ia input is necessary for recruiting high-threshold motor units, enhanced cutaneous stimulation might indirectly affect alpha motor neuron activation(11). Based on this mechanism, some reports concluded that a decline in alpha motor neuron activity due to AMI could be partially restored by cutaneous stimuli(15, 16).

Previous studies used maximal effort strength during knee extension exercise to evaluate the effect of cutaneous stimulation on AMI(15, 16). However, motor function during coordinated functional movements should also be examined, as these movements reflect the actual state of the human body more closely than a simple movement like knee extension. Moreover, in actual situations, especially during sports activities, coordinated functional movements cannot always be accomplished as predicted because of various perturbations. Unpredictable events could lead to undesirable outcomes, such as falling. Therefore, it is important to clarify how the neural system adapts to unpredictable tasks incorporated into coordinated functional movements. Since landing on a single leg is one of the most common tasks performed during coordinated functional movements such as walking, running, and jumping, a study design using unpredictable tasks involving single leg landing will be widely applicable to various types of human movements. To conduct a study aimed at characterizing the effect of unpredictable tasks on motor control, simulating coordinated functional movements under unpredictable conditions will be necessary. To date, no study has investigated the effect of AMI on coordinated functional movements under unpredictable conditions.

Therefore, the purpose of the present study was to investigate the effect of tactile stimulation on reflex changes induced by AMI during unpredictable landing performances. However, it is impossible to determine the direct effect of AMI due to actual injury on reflexive motor control during unpredictable landing performance, because pre- and post-injury reflex changes cannot be determined. Thus, in the present study, the simulated AMI in which neurological changes are induced by prolonged vibration stimulation(15, 16) was used, and data obtained from the same subjects before and after induction of neurological changes similar to AMI were compared.

## Methods

### Participants

Informed consent was obtained from all participants before participating in the study. All procedures conformed with the Helsinki Declaration and were approved by the Committee on Human Experimentation at the National Defense Academy of Japan. Participants were 20 healthy volunteers (13 males and 7 females; age: 22.2±6.9 years, height: 168.1 ±4.8 cm, weight: 61.6±5.4 kg, mean ± SD) who had no history of major injury in the lower limbs. All participants were recruited between August 2020 and March 2021.

### Experimental procedure

To warm up, all participants pedaled on an exercycle for five minutes. Participants performed single leg landing on one limb (randomly selected) after receiving instructions on the landing technique and practicing landing from 15 cm and 30 cm heights, as well as surprise landing, to familiarize themselves with unilateral landing tasks. All participants performed six types of unilateral landing tasks (Tasks 1 to 6; 15 cm normal landing (15NL), 30 cm normal landing (30NL), surprise landing (SL), 30 cm normal landing following vibration (30NLV), surprise landing following vibration (SLV), and surprise landing following vibration with Kinesiology taping (SLK). After completing Tasks 1 to 3 in that order, vibration stimulation was applied. Immediately after the application of vibration stimulation, participants performed 30NLV (Task 4) five times, followed by SLV (Task 5) and SLK (Task 6) in a random order.

### Vibration protocol

Vibration stimulation was applied manually using the Hit Masser (Kinesio Co., Tokyo) to the infrapatellar tendon as previously described(2, 15, 16). The simulated AMI using prolonged vibration stimulation to induce reflex changes during unpredictable landing performances would be affected by tactile cutaneous stimulation. Thus, it is essential to determine whether or not the effect of the simulated AMI on motor control during unpredictable landing would induce changes in motor control, as it is otherwise impossible to address the purpose of the present study. The advantage of using the simulated AMI is that we could focus on the effect of neurological changes, so that other factors such as pain and pathological conditions could be excluded (because neurological changes are induced by vibration stimulation in the simulated model). Moreover, it allowed us to compare data obtained from the same subjects before and after induction of neurological changes, as these changes were induced intentionally by means of prolonged vibration stimulation.

### Landing task procedures

Participants wearing shoes were instructed to step off the take-off box and land on a cross marked on the landing surface, 40 cm in front of them. For 30NL, participants stepped off the box and landed directly on the floor; for 15NL, a 15 cm landing platform was placed in front of the 30 cm take-off box (Fig.1 A). Participants performed 15NL and 30 NL five times each, followed by the surprise landing session. The solid 15 cm landing platform was removed and replaced by one with a false floor. As the surface of this floor was made of a very light semi-rigid corrugate board, which dislodged easily under load, participants unexpectedly fell through the false floor to the solid floor 15 cm below (Fig.1 B). The surprise landing session consisted of 15NL and SL trials: the total number of landing tasks performed ranged from eight to 10 and included one to three SL and five to nine 15NL trials. Participants were not notified which trials were SL or 15NL, and performed the trials for a random number of times (either 1–3 times or 8–10 times). Moreover, appearances of the normal and false landing surfaces were indistinguishable, and since participants were asked to move to a place where they could not see the setting after each trial, they could not figure out if the landing surface had been changed. Finally, vibration stimulation was applied for 20 minutes, and participants performed 30NL five times and SLV one to three times (a total number of landing tasks was set to four to six).

**FIGURE 1.**
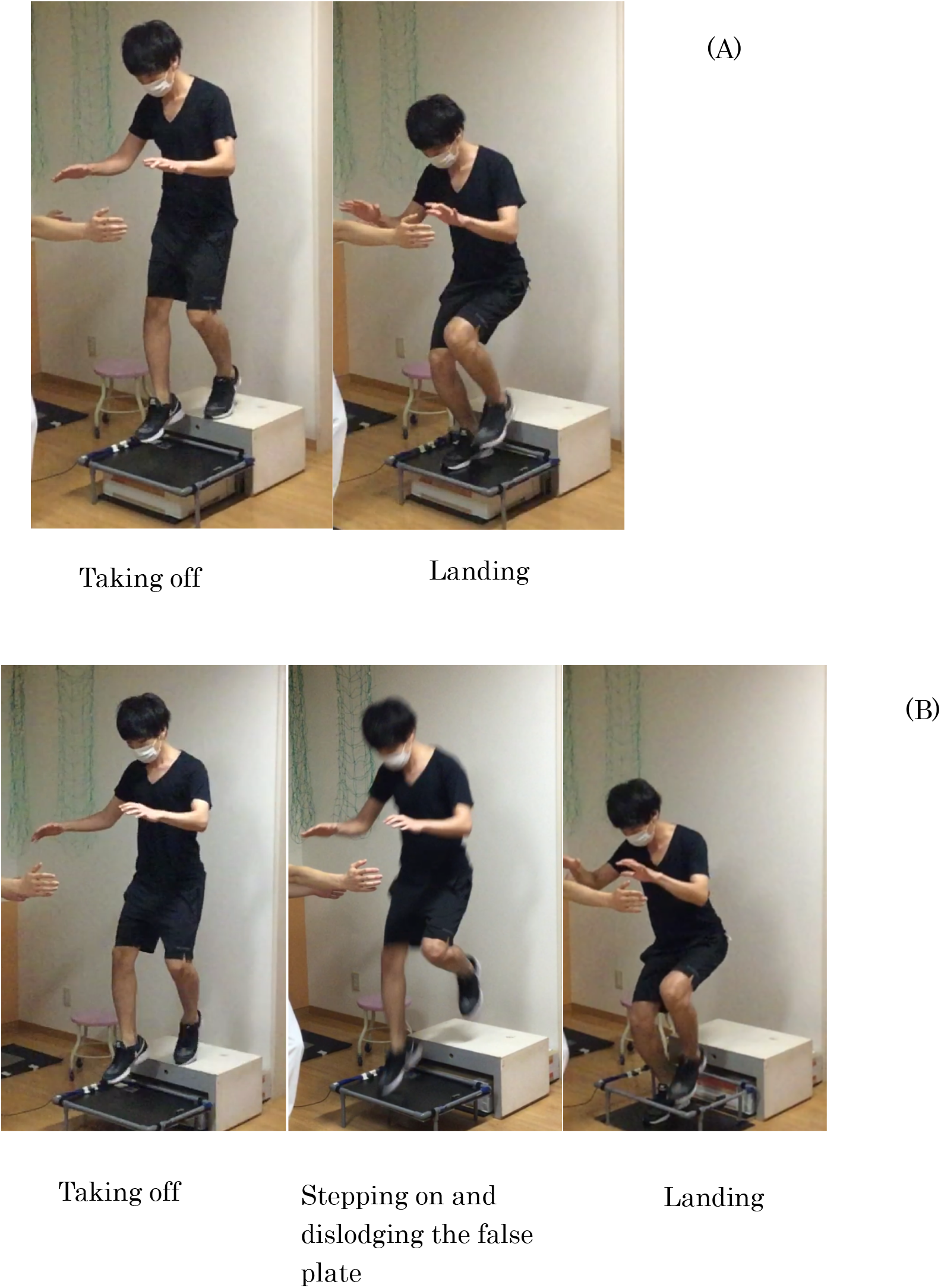
(A) 15 cm normal landing. A solid 15 cm landing platform was placed in front of a 30 cm take-off box. (B) Surprise landing. As the surface of the floor dislodged easily under load, participants unexpectedly fell through the false landing surface to the solid floor 15 cm

Electronic contact mats attached to the upper surface of the 30 cm take-off box, 15 cm landing platform, and the floor in the landing area provided the timing of when participants left the box and the moment of contact with the landing surface. A photocell provided data on the timing of contact with the false floor. Signals from these devices were transmitted to Signal 5.11 (CED, Ltd., Cambridge, UK) via an A/D conversion unit (Micro 1401, CED, Ltd., Cambridge, UK) at 2 kHz. The procedures in the present study were modified from those used in a previous study with the same surprise landing protocol(17).

### Mean percentage change in electromyography (EMG) normalized by maximal EMG

EMG signals from the vastus lateralis (VL), vastus medialis (VM), and lateral hamstrings (LH) were obtained. The skin was prepared by shaving, abrading, and cleaning the area before placing bipolar electrodes (inter-electrode distance: 20 mm) over these muscles according to SENIAM guidelines(18). The electrodes were connected to a measurement unit (SX230-1000, Biometrics, Ltd., Gwent, UK) that filtered data between 5–450 Hz. EMG data were transmitted to Signal 5.11 (CED, Ltd., Cambridge, UK) via an A/D conversion unit (Micro 1401, CED, Ltd., Cambridge, UK) at 2 kHz. EMG data were collected from the time of leaving the box to roughly 6000 ms post-landing. For 30NL and 30NLV, the averaged waveform of collected EMG data was obtained. For surprise landing tasks (SL, SLV, and SLK), EMG data from the first trials were used in analysis. Averaged waveforms for 30NL and 30NLV and data from the first trials for SL, SLV, and SLK were used as variables.

To allow for comparisons across participants, EMG signals were normalized to EMG amplitude during isometric maximal voluntary contraction (MVC). To obtain EMG signals normalized by maximal EMG during maximal effort quadriceps and hamstring contractions, these tasks were performed with the knee flexed at 90 degrees. Two maximal efforts of approximately five seconds were recorded, and the highest EMG amplitude across a one second moving average was used.

### Knee joint angular velocity

Electrogoniometers (DL-260, S&ME, Ltd., Tokyo, Japan) were used to obtain data on knee angles and joint angular velocity during landing tasks. They were attached to the skin on either the lateral or medial aspect of the knee depending on the alignment of the thigh and leg. Signals were transmitted to Signal 5.11 (CED, Ltd., Cambridge, UK) via an A/D conversion unit (Micro 1401, CED, Ltd., Cambridge, UK) at 2 kHz. Maximal flexion angle following touch-down on the ground was obtained and divided by the time from touch-down to when maximal flexion angle was reached to obtain the angular velocity of the knee joint (deg/sec). For 30NL, the averaged waveform of collected goniometry data was obtained. For SL, SLV, and SLK, goniometry data from the first trials were used in the analysis.

### Estimated vertical ground reaction force

An accelerometer (4Assist Inc., Tokyo, Japan) was used to obtain data on acceleration during landing tasks. The accelerometer was placed on the top edge of the sacrum of each participant. Signals were transmitted to Signal 5.11 (CED, Ltd., Cambridge, UK) via an A/D conversion unit (Micro 1401, CED, Ltd., Cambridge, UK) at 2 kHz. Vertical acceleration data of each participant were multiplied by their body mass to obtain the reaction force from the ground. Maximal vertical acceleration data following touch-down on the ground were obtained and divided by the time from touch-down to when maximal amplitude was reached to obtain the change rate of the data. For 30NL, the averaged waveform of collected estimated ground reaction force data was obtained. For SL, SLV, and SLK, estimated ground reaction force data from the first trials were used in analysis.

### Tactile stimulation to the skin around the knee

Kinesiology tape (Nitto Denko Corp., Tokyo, Japan), which is commonly used for clinical purposes, was applied to approximately 3 cm above the patella to the tibial tuberosity to cover each participant’s knee joint. The tape was applied immediately after vibration stimulation and left on during MVC measurements. To reduce the placebo effect, participants were not notified about the advertised effects of the tape. The detailed method of tactile stimulation application has been described previously(15).

### Data analyses

All data were processed and analyzed offline using Signal 5.11 (CED, Ltd., Cambridge, UK). For 30NL, EMG signals were full-wave rectified, and average EMG amplitude (normalized to % of MVC) was examined across different epochs associated with landing tasks and averaged across all trials. Three epochs were chosen relative to the time of landing (0 ms): −60 to 0 ms, 31 to 60 ms, and 61 to 90 ms. These epochs were chosen to provide measurements of pre-landing EMG activity in anticipation of landing (−60 to 0 ms), as well as two post-landing EMG activities that include short latency (31 to 60 ms) and medium latency (61 to 90 ms) stretch reflex responses, as reported previously(19, 20). For SL, given that our focus was to compare post-landing stretch reflex responses in SL and 30NL, EMG signals were analyzed for the post-landing short and medium latency epochs only. Neither pre-landing nor post-landing EMG amplitude was analyzed for 15NL, as this condition was only used to familiarize the participants with landing performance.

### Statistical analyses

All data were analyzed using R (IBM Corporation, Armonk, USA). Data were initially assessed by Q-Q plots to examine the normality of the sample distribution, and hence, to determine the use of non-parametric inferential statistical tests. As mentioned, the purpose of the present study was to investigate how neurological changes during unpredictable landing would be affected by cutaneous stimulation. Since it was unclear if prolonged vibration stimulation used in the simulated AMI actually affected motor control during unpredictable landing, it was essential to first examine the effect of the simulated AMI. To this end, the Wilcoxon signed-rank test was used to compare the following variables: average EMG amplitude for each muscle, average knee joint rotation velocity, and magnitude of estimated vertical ground reaction force between SL and SLV. In order to address the purpose, SLV and SLK data were compared using the Wilcoxon signed-rank test. Average EMG variables for the short latency (31-60 ms) and medium latency (61-90 ms) epochs were analyzed separately. For all analyses, the alpha level was set to 0.05. In addition, to confirm that the SL protocol properly reproduced unpredictable conditions, average EMG amplitudes for each muscle were compared between 30NL and SL using the Wilcoxon signed-rank test.

## Results

### Post-landing short latency EMG (31 to 60 ms) (Table 1A)

For all muscles, the Wilcoxon signed-rank test revealed no change in the average amplitude of EMG signals during the short latency epoch between SLV and SL (Table 1A). In addition, there was no change in the average amplitude of EMG signals during the short latency epoch between SLK and SLV (Table 1A).

**Table 1.**
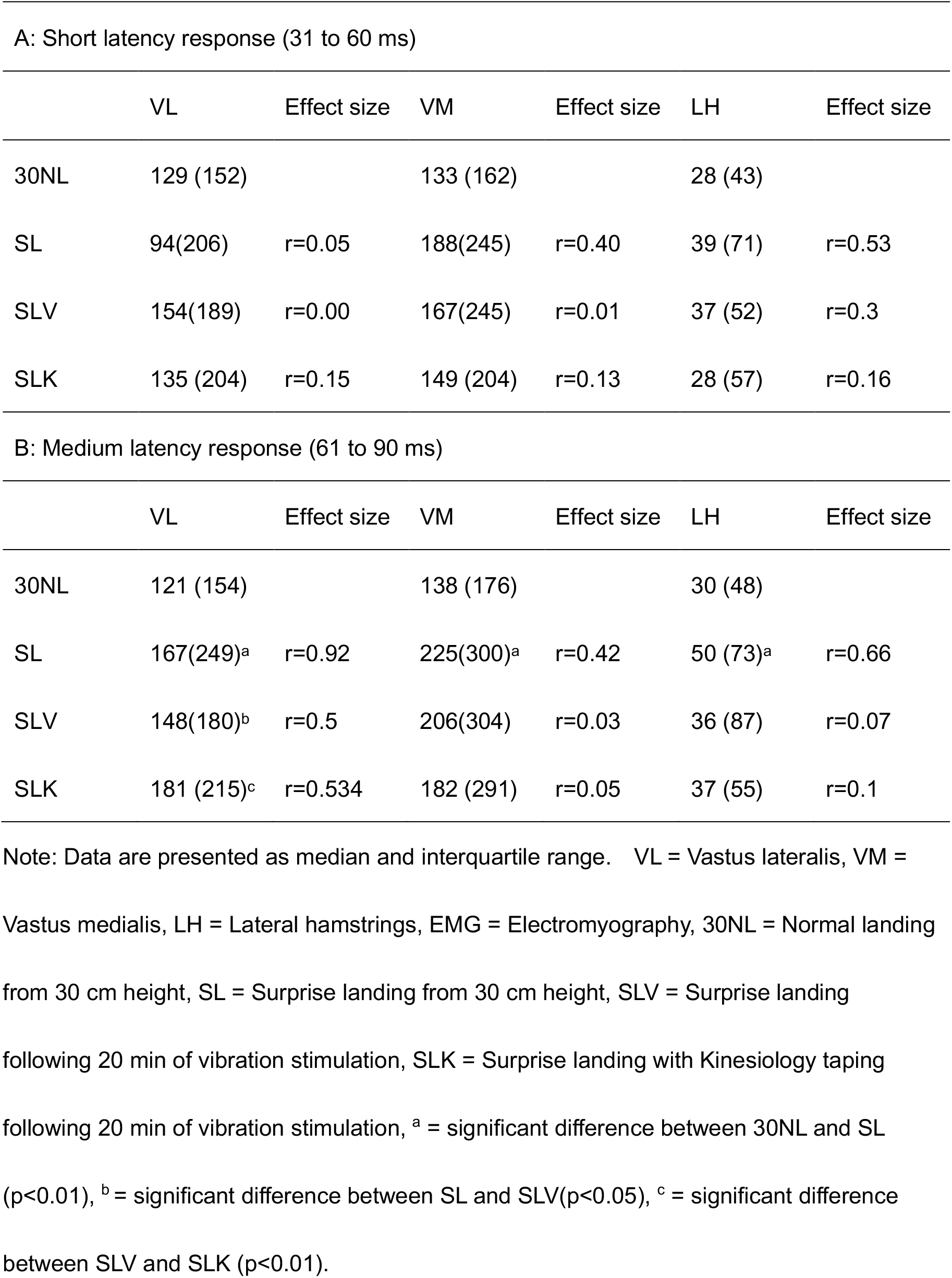
Average amplitude of EMG signals from the vastus lateralis, vastus medialis, and lateral hamstrings.

### Post-landing medium latency EMG (61 to 90 ms) (Table 1B)

The Wilcoxon signed-rank test revealed that the average amplitude of EMG signals from the VL during the medium latency epoch in SLV was significantly smaller than that in SL (p<0.01) (Table 1B). No significant differences were observed for the VM and LH.

The Wilcoxon signed-rank test also revealed that the average amplitude of EMG signals from the VL during the medium latency epoch in SLK was significantly larger than that in SLV (p<0.01) (Table 1B). No significant differences were observed for the VL, VM, and LH.

### Knee joint angular velocity (Table 2)

The Wilcoxon signed-rank test did not revealed no change between between SLV and SL (SLV: 275±69 deg/sec, SL: 326±91deg/sec), and between SLK and SLV (SLK: 317±87 deg/sec, SLK: 317±87 deg/sec).

**Table 2.** Median rate of force development caluculated from estimated vertical ground reaction force obtained from acceleration data and knee joint angular velocity.

|  | EGRF (N/s) | Effect size | KAV (deg/sec) | Effect size |
| --- | --- | --- | --- | --- |
| 30NL | 13091 (13150) |  | 277 (290) |  |
| SL | 40538 (41688)<sup>a</sup> | r=0.90 | 317(366) | r=0.41 |
| SLV | 30239 (30494) | r=0.40 | 324(382) | r=0.15 |
| SLK | 32702 (33819) | r=0.11 | 310 (361) | r=0.24 |

Note: Data are presented as median and interquartile range. EGRF = Estimated vertical ground reaction force, KAV = Knee joint angular velocity, 30NL = Normal landing from 30 cm height, SL = Surprise landing from 30 cm height, SLV = Surprise landing following 20 min of vibration stimulation, SLK = Surprise landing with Kinesiology taping following 20 min of vibration stimulation. <sup>a</sup> = significant difference between 30NL and SL (p<0.01).

### Rate of force development caluculated from estimated vertical ground reaction force obtained from acceleration data (Table 2)

Acceleration data were multiplied by body mass to obtain the estimated vertical reaction force from the ground and divided by the time to reach maximal value from base line data. The Wilcoxon signed-rank test did not revealed no change between between SLV and SL (SLV: 30770±30291N/sec, SL: 326±91deg411859±299130 N/sec), and between SLK and SLV (SLK: (33634±33935N/sec, 30770.31731±30291N/sec).

## Discussion

In the present study, vibration stimulation (50 Hz) was applied to the infrapatellar tendon for 20 minutes. This stimulation temporally diminished the activity of alpha motor neurons while exerting MVC. Previous studies have demonstrated that prolonged vibration stimulation to muscles hinder the recruitment of high-threshold motor units as a result of attenuation of Ia afferent feedback in healthy subjects(14). Indeed, several studies have reported a decline in the activity of alpha motor neurons upon prolonged vibration stimulation in healthy subjects(2, 14, 7). This hindrance to the recruitment of motor units was also observed in patients suffering from persistent quadriceps femoris weakness(2, 4, 3). Muscle weakness observed after prolonged vibration stimulation might have a neurophysiological underpinning similar to that observed in those patients. Previous studies using the simulated AMI with prolonged vibration stimulation reported that cutaneous stimulation such as tactile and electrical stimulation to the skin around the knee joint prevented the decrease in both strength and EMG activity(15, 16). Thus, the attenuated alpha motor neuron activity due to prolonged vibration stimulation might be partially rescued by cutaneous stimuli. In the present study, we examined landing movements following neurological changes using the simulated AMI (i.e., induced by vibration stimulation) and found that EMG signals from the VL during the medium latency epoch were significantly smaller in SLV than in SL. Unpredictable landing normally leads to a gain of medium latency stretch reflex(17, 21), and in this regard, the results of the present study were consistent with previous reports(17, 21).

The gain of medium latency stretch reflex in the VL, which is reportedly observed during unexpected landing, was not observed after the application of prolonged vibration. Given the similarity between the neurological changes induced by prolonged vibration stimulation and those caused by AMI, our results provide evidence that AMI hinders the gain of reflex responses during unpredictable landing in the VL, which are normal responses in healthy individuals. Since no change was observed in the hamstrings, AMI may not affect the function of hamstrings.

In the present study, tactile stimulation was applied following the induction of neurological changes using the simulated AMI during unpredictable landing. Consequently, tactile stimulation restored the hindered reflex gain in the VL. With regard to the relationship between sensory input (e.g., tactile stimulation) and motor neuron activity, sensory input from the skin around the joint could activate gamma motor neurons to modulate Ia afferent activity, and the same sensory input could also indirectly activate alpha motor neurons(11). The gain of afferent feedback due to sensory input from the skin might be the mechanism of how tactile stimulation restores the declined reflex in the simulated AMI.

A previous study investigating the effect of unpredictable landing on reflex patterns in the lower limb muscles of patients with ACL rupture who were suffered from AMI found a gain of medium latency reflex in the VL(17). Our results were not consistent with this report, possibly because patients with ACL rupture had already acquired proper skills to adapt to unpredictable events prior to testing, since they were potential copers who could adapt to strenuous activities without reconstruction(22). Indeed, all those patients who participated in the previous study met the following criteria of screening examinations: (a) a limb symmetry score >80% in the 6-m timed hop test, (b) Activities of Daily Living Scale score >80%, (c) global rating of knee function >60%, (d) no more than one episode of giving way, and (e) no pain, no effusion, and full range of motion. In contrast, it is likely that participants of the present study had not adapted to unpredictable events, i.e., they were required to perform unpredictable landing soon after vibration stimulation was applied(17). Given the discrepancy in response to neural changes associated with AMI between the present and previous studies, the gain of medium latency reflex in the VL might be an essential factor that clinicians should focus on during rehabilitation.

## Conclusions

The use of the simulated AMI allowed us to focus on the potential effect of AMI itself, since neurological changes similar to those caused by AMI could be induced by using prolonged vibration stimulation. Our findings suggest that AMI could alter motor control patterns during unpredictable landing, and that tactile stimulation could restore the altered motor control to a normal state. These results support the usefulness of tactile stimulation for maintaining normal motor control patterns in injured individuals.

## Acknowledgements

This work was supported by JSPS KAKENHI Grant Number JP 20K09515.

